# HIV-1 interactions with sialic acid-binding bacterial lectins promote virus infectivity in vitro and mucosal transmission in humanized mice

**DOI:** 10.64898/2026.05.05.722898

**Authors:** Clauvis Kunkeng Yengo, Xiaomei Liu, Ries J. Langley, Frida Avila, Manish Sagar, Christina Ochsenbauer, Barbara A. Bensing, Catarina E. Hioe

## Abstract

Most HIV-1 transmission occurs at mucosal surfaces, which are colonized by the host microbiota. However, interactions between HIV and bacteria or bacterial products derived from the human microbiome are poorly characterized, and their biological consequences are largely unexplored. Here, we evaluated the effects of sialic acid-binding lectins expressed by bacterial species ubiquitous in the human microbiota on HIV-1 infectivity using viruses produced in 293T cells and human primary cells. We demonstrated that these bacterial lectins enhanced HIV-1 infectivity in a sialoglycan-dependent manner. Specifically, Siglec-like binding region lectins (SLBR-N, SLBR-H, and SLBR-B) from *Streptococcus gordonii* and Staphylococcal superantigen-like lectins (SSL3, SSL4, and SSL11) from *Staphylococcus aureus* increased HIV-1 infectivity to varying extents, depending on lectin type and virus strain. Among these lectins, SLBR-N exhibited the greatest potency, corresponding with its superior ability to bind virions and promote virus-cell attachment. This enhancing activity was observed for direct infection of TZM-bl reporter cells and primary CD4+ T cells, as well as trans-infection in the presence or absence of the mannose-binding host lectin DC-SIGN. Importantly, these findings were corroborated in vivo using humanized mice, in which pre-exposure to SLBR-N promoted rectal HIV-1 transmission and increased viral burdens in plasma and splenic cells. Collectively, the data show sialoglycan-binding bacterial lectins as microbial factors that can enhance HIV-1 transmission at mucosal surfaces, highlighting a potential direct role for the microbiota in modulating HIV-1 acquisition risk.

**Author Summary:** HIV is commonly transmitted from one person to another across mucosal surfaces, such as those lining the genital and rectal tracts, which are densely populated by bacteria that make up the human microbiota. Yet, surprisingly little is known about how these bacteria and the molecules they produce influence HIV infection. In this study, we investigated a group of bacterial proteins known as sialic acid-binding lectins that are expressed by common members of the human microbiome: Siglec-like binding region lectins from *Streptococcus gordonii* and superantigen-like lectins from *Staphylococcus aureus*. Using multiple HIV strains and several types of target cells, we demonstrate that lectin binding to HIV can increase virus attachment to target cells and thereby enhance infection, although the magnitude of this effect varies among lectins and virus strains. Lectin binding also facilitates HIV spread from cell to cell and promotes mucosal HIV infection in a humanized mouse model, resulting in a higher viral burden in the blood and tissues. These findings identify bacterial lectins as important factors that can influence HIV infection and implicate a potential role for the human microbiota in determining susceptibility to HIV infection.

## Introduction

Most human immunodeficiency virus type 1 (HIV-1) transmission occurs through the mucosa during sexual contact, perinatal events, and breastfeeding (1, 2). At the mucosal surfaces, the virus interacts not only with host cells but also with bacteria in the microbiota. Indeed, bacterial dysbiosis is linked with increased risk of HIV-1 acquisition (3–6), although this has been attributed mainly to the associated inflammation and compromised mucosal barrier.

Mucosal microbial communities are dynamic and modulated by hormonal changes and contraceptive use, among other factors (7–12). Depot-medroxyprogesterone acetate (DMPA or Depo-Provera), an injectable contraceptive that has been associated with increased HIV-1 acquisition risk (13–17), induces shifts in the vaginal microbiota, signified by reduced Lactobacillus dominance, enrichment of vaginosis-associated bacteria such as Prevotella, and retention of a few community networks that include Streptococcus and Mycoplasma (18–20). Despite these associations, direct mechanistic links between microbiome changes and HIV-1 transmission remain poorly defined.

Among vaginosis-associated bacteria, *Prevotella timonensis* encodes mucus-degrading enzymes, including fucosidases and neuraminidases (sialidases) that degrade mucin O-glycans, which are critical for maintaining healthy cervicovaginal barriers (21, 22). *P. timonensis* also enhances HIV-1 infection in vaginal CD4+ T cells and promotes trans-infection from CD1c+ dendritic cells and vaginal Langerhans cells to CD4+ target cells (23–25). Notably, these effects are driven by bacteria-cell interactions (26), rather than direct bacteria-virus engagement.

Investigations on direct HIV-1-bacteria interactions have been limited, although intimate connections between bacteria and other viruses have been well documented. Human norovirus binds the histo-blood group antigen (HBGA) glycans from commensal *Enterobacter cloacae* to infect B cells (27–29). Poliovirus binds bacterial lipopolysaccharide (LPS) via its capsid, increasing virus thermostability, resistance to bleach, and virus replication in the intestine (30, 31). Similarly, murine mammary tumor virus (MMTV) binds bacterial LPS via LPS-binding host proteins (CD14, MD2, and TLR4) incorporated in the virus envelope (32), and MMTV-bound LPS triggers TLR4 to induce IL-6 and IL-10, thereby creating an immunosuppressive environment that permits virus persistence (33, 34).

At the mucosal sites such as the urogenital tract, oral cavity, and gastrointestinal tract, commensal and pathobiont bacteria express a variety of lectins that recognize glycans on host cells, mucus, and extracellular matrix proteins, and play key roles in bacteria adhesion, biofilm formation, and immune evasion (35). We previously revealed that bacterial lectins also interact with HIV-1 virions and the virus envelope glycoprotein (Env) in a glycan-dependent manner resulting in distinct outcomes depending on the lectins. HIV-1 binding to sialylated O-glycan-specific SLBR lectins from *Streptococcus gordonii* resulted in enhanced virus infectivity, whereas the interaction with high-mannose N-glycan-specific FimH lectin from *E. coli* caused no changes (36), implicating the role of specific bacterial lectins as modulators of HIV-1 infection. However, the previous study was limited to these lectins and a few virus strains, and the biologic consequences in vivo were unexplored.

The present study investigated sialoglycan-binding lectins from *S. gordonii* and *Staphylococcus aureus* for the ability to augment infection across HIV-1 strains, including viruses produced in primary human CD4 T cells. *S. gordonii* is a commensal bacterial species that is found in the oral cavity, colonizing tooth surfaces beginning in infancy and is linked to tooth decay and endocarditis (37–39). In addition, *S. gordonii* can also be found in the skin, upper respiratory tract, and intestine (39). *S. aureus* is a pathobiont that colonizes humans soon after birth. *S. aureus* is ubiquitous, present in 30-50% of the human population, including both healthy and hospitalized individuals, with reservoirs in the nasopharynx, the skin, the genital tract, and the intestinal tract (40–42). Glycan-dependent HIV-1 interactions with these bacterial lectins were ascertained using a glycan competition assay and lectin mutants. In addition to direct infection, we evaluated the effect of bacterial lectins on HIV-1 trans-infection in the presence and absence of the host lectin, DC-SIGN. Finally, we assessed lectin activity on rectal HIV-1 infection using a humanized mouse model. Data from these experiments highlight the important role of sialic acid-binding lectins from common bacteria in the human microbiota in influencing HIV-1 transmission.

## Results

### Streptococcal SLBR lectin-mediated enhancement of HIV-1 infectivity varies by viral strain and lectin specificity

We previously showed that Siglec-like binding region (SLBR) lectins from *Streptococcus gordonii* enhanced virus infectivity through interactions with glycan structures on HIV virions (36). However, the previous experiments were performed in the presence of a polycation, diethylaminoethyl (DEAE) dextran, which augments HIV infection by countering the electrostatic-repulsive forces between virus and cell surface (43–45). Under those conditions, only modest 2- to 4-fold increases in infectivity were observed (36). To isolate the effects of bacterial lectins, DEAE dextran was excluded from experiments in the present study.

In addition to SLBR lectins, we also evaluated staphylococcal superantigen-like (SSL) lectins from *Staphylococcus aureus*. Although both SLBR and SSL lectins recognize terminal sialic acids, they exhibit distinct glycan preferences (**Figure 1**). SLBR-N preferentially binds terminal α2,3 sialic acids present in 3’ sialyllactosamine (sLn) and sialyl Lewis X (sLe^X^) on core 2 *O*-glycans (46, 47). SLBR-B recognizes sialyl T-antigen on core 1 *O*-glycans, whereas SLBR-H has broader specificity, binding sLn and other sialoglycans on both core 1 and core 2 *O*-glycans (38, 48, 49). SSL lectins similarly recognize the minimal core ligand sLn but show greater affinity for sLe^X^ (50–53), although each has a preference for terminal α2,3 sialic acids on distinct N-linked or O-linked oligosaccharides (54).

**Figure 1.**
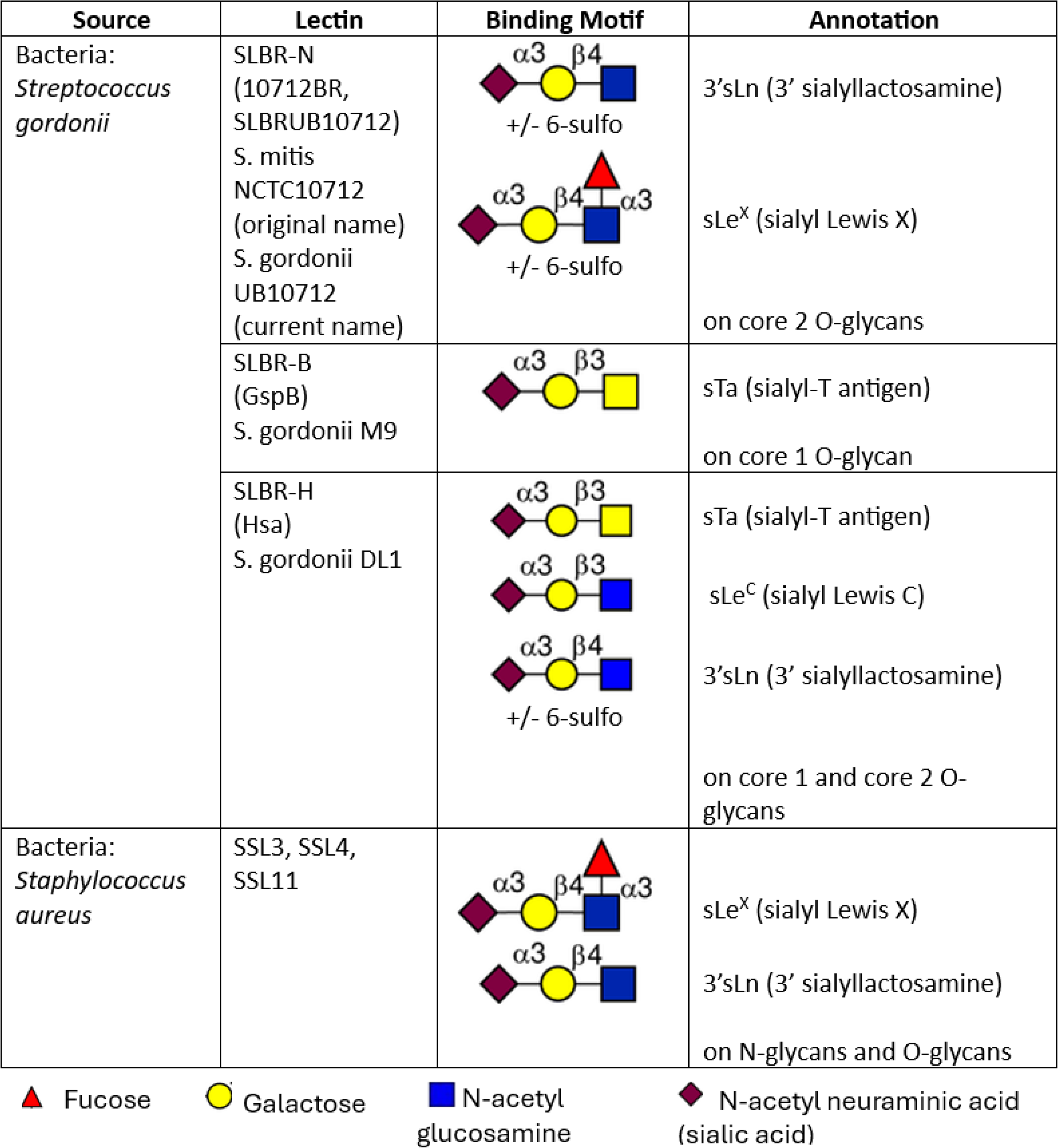
Glycan specificities of SLBRs and SSLs. Glycan structures recognized by each bacterial lectin were previously reported (38, 48, 49, 50–53).

To assess SLBR-mediated enhancement of HIV infectivity, we initially tested HIV-1 infectious molecular clones (IMCs) JRFL (chronic) and REJO.c (transmitter/founder) (55), produced from 293T cells. IMC virions were pre-incubated with titrated concentrations of individual lectins at 37°C for 1 hour and then added to TZM-bl reporter cells. Infectivity was quantified after 48 hours. Lectin treatment enhanced virus infectivity in a dose-dependent manner, with the magnitude of enhancement varying by both viral strain and lectin (**Figure 2A**). Comparable results were observed whether the lectins were pre-incubated with virus or added simultaneously to virus and cells (**Supplemental Figure S1**). In contrast, we previously showed that SLBR-N had no effect when it was added to TZM-bl target cells before addition of virus or after infection, indicating that this lectin exerts its activity on virions and not on target cells (36). Notably, SLBR-N produced greater enhancement than SLBR-H or SLBR-B for both JRFL and REJO.c (**Figure 2A**). These results are also consistent with our prior findings using other HIV-1 strains, including clade C Z331M and CRF01_AE CMU06 (36).

**Figure 2.**
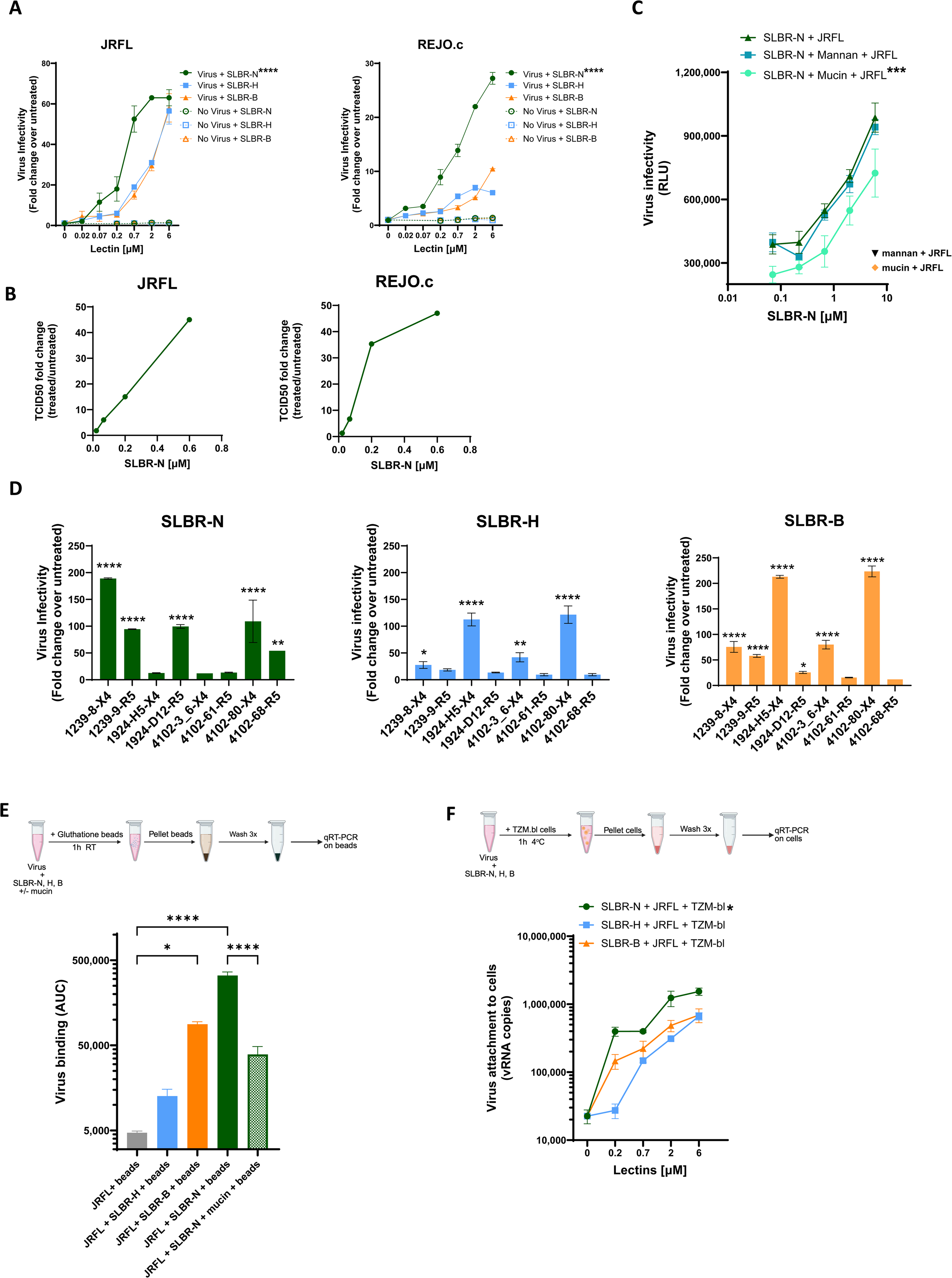
SLBR-N, SLBR-H, and SLBR-B enhance virus infectivity to varying levels across HIV-1 strains. **(A)** Effects of SLBR lectins on the infectivity of HIV-1 virions produced in 293T cells. JRFL and REJO.c IMC virions were pre-incubated with titrated amounts of lectins at 37 °C for 1 hour, and infectivity in TZM-bl reporter cells was quantified by beta-galactosidase activity after 48 hours. Fold changes over untreated virus across titrated lectin concentrations are shown. Effects of lectins alone, without virus, on TZM-bl cells are also shown. Mean and SD are presented. Infection with untreated JRFL and REJO.c yielded 8,899 RLU and 15,973 RLU, with background (uninfected cells) of 4,164 RLU and 4,172 RLU, respectively. RLU: relative luminescence unit. **** p<0.001 for SLBR-N vs. SLBR-H and SLBR-B by two-way ANOVA with Šídák’s multiple comparison. **(B)** Enhancing effect of SLBR-N on HIV-1 infectivity as measured by TCID50. JRFL and REJO.c IMC virions were serially diluted and incubated with or without SLBR-N (0.6 to 0.02 µM) for 1 hour at 37°C. Reciprocal virus dilutions required to reach 50% infection (TCID₅₀) was calculated in GraphPad Prism using a non-linear fit model. Fold changes in TCID₅₀ of untreated vs. SLBR-N-treated virus at each lectin concentration tested are shown. **(C)** Enhancing activity of SLBR-N was decreased by O-glycan-bearing mucin but not high-mannose N-glycan-bearing mannan. SLBR-N was pre-incubated with 1 mg/mL mucin or mannan at 37 °C for 1 hour, followed by incubation with JRFL virions for an additional 1 hour at 37 °C and then addition of TZM-bl cells. *** p<0.001 for comparing JRFL + SLBR-N treated with mucin vs. JRFL + untreated SLBR-N by two-way ANOVA. p >0.05 for JRFL + SLBR-N treated with mannan vs. JRFL + untreated SLBR-N. Orange diamond and black triangle: RLU for JRFL infection in the presence of only mucin or mannan, respectively. **(D)** Varying effects of SLBR-N lectins on HIV-1 strains produced in primary CD4 T cells. Clade B R5-tropic or X4-tropic IMCs were produced in activated CD4 T cells from the same PBMC donor. Virions were treated with SLBR lectins and virus infectivity was measured as in panel A. Fold changes over untreated control at the highest lectin concentration are shown. ****p<0.0001 **p<0.01 *p<0.05 vs. no lectin by ANOVA. p≥0.05 is left unmarked. **(E)** Different levels of virus binding to SLBR-N, SLBR-H, and SLBR-B lectins. JRFL virions were pretreated with each GST-tagged SLBR lectin and then incubated with glutathione beads, and the amount of virus capture on beads was measured by RT-qPCR. The blocking activity of mucin was assessed to verify O-glycan-dependent virus capture by SLBR-N. ****p<0.0001 *p<0.05 by ANOVA with Tukey’s multiple comparisons test. p≥0.05 is left unmarked. **(F)** Varying levels of virus attachment to TZM-bl target cells in the presence of SLBR-N, SLBR-H, and SLBR-B lectins. Virus was pre-treated with each lectin for 1 hour at 37°C and then incubated with TZM-bl cells for 1 hour at 4°C. The amount of cell-associated virus was quantified by RT-qPCR. * p<0.05 for SLBR-N vs. SLBR-H and SLBR-B by two-way ANOVA with Tukey’s multiple comparison. Mean ± SD are presented in panels A, C, D, E and F; data from one experiment out of two or more repeat experiments are shown.

We subsequently measured the effect of SLBR-N on virus infectivity as measured by 50% tissue culture infectious dose (TCID50). JRFL and REJO.c IMCs were titrated, treated with SLBR-N (0.02 to 0.6 µM) for 1 hour at 37°C, and then added to TZM-bl cells. SLBR-N increased virus dilutions required for 50% infection of both virus strains in a dose-dependent manner (**Figure 2B, Supplemental Figure S2**), confirming that this lectin indeed increased virus infectivity.

To evaluate the contribution of specific glycan-lectin interactions, we assessed the effects of mucin (Type I-S), which is richly decorated with sialylated O-glycans, and mannan, which contains high-mannose N-glycans. Mucin treatment markedly reduced SLBR-N-mediated enhancement of infectivity, whereas mannan had no discernible effect (**Figure 2C**). Neither mucin nor mannan alone altered baseline virus infectivity. These results demonstrate the role of sialylated O-glycans in SLBR-N-mediated enhancement of HIV-1 infectivity.

Because host producer cells influence virus glycan composition (56–58), we next examined SLBR activity on HIV-1 virions produced in primary CD4 T cells. Eight clade B IMCs, which express *in cis* heterologous Env with X4 or R5 tropism, were propagated in activated PBMCs from a single donor. Similar to viruses produced in 293T cells, SLBR treatment enhanced infectivity of PBMC-derived viruses, although the magnitudes of enhancement varied substantially among strains (**Figure 2D**). Astonishingly, the pattern of enhancement differed by lectin. For example, infectivity of 1239-8 (X4) was most strongly enhanced by SLBR-N, followed by SLBR-B and SLBR-H, whereas 1924-H5 (X4) was unaffected by SLBR-N but showed increased infectivity in the presence of SLBR-B and SLBR-H. Interestingly, SLBR-H significantly enhanced infectivity of only X4-tropic strains and not R5-tropic strains, while SLBR-N and SLBR-B displayed activity on both X4- and R5-tropic strains. Nonetheless, all eight strains tested exhibited significant enhancement by at least one SLBR lectin. These data demonstrate differential sensitivity of virus strains to lectin-mediated enhancement of infectivity, indicating substantial heterogeneity in the abundance and composition of core1 versus core2 O-linked sialoglycans among HIV-1 strains.

### SLBR lectins augment HIV infectivity by binding virions and promoting virus-cell attachment

To determine whether differences in infectivity enhancement reflected variations in lectin-virion interactions, we performed virus capture assays using Glutathione S-Transferase (GST)-tagged SLBR lectins. JRFL virions were pre-incubated with individual SLBR lectins and then captured by glutathione beads. Following extensive washing, lectin-bound virions captured on the beads were quantified by qRT-PCR. JRFL virions were captured most efficiently in the presence of SLBR-N, followed by SLBR-B and SLBR-H (**Figure 2E**), demonstrating that SLBR-N has greater binding capacity for JRFL than the other two lectins. Treatment with sialyated O-glycan-rich mucin significantly reduced SLBR-N-mediated virus capture, further highlighting the specific contribution of sialoglycans to HIV-1 interactions with SLBR-N.

We further examined whether lectin binding to virions facilitated virus attachment to cells. Lectin-treated JRFL virions were incubated with TZM-bl cells at 4°C for 1 hour to allow virus attachment without entry. After extensive washing, cell-associated viral RNA (vRNA) was quantified by qRT-PCR. Lectin treatment increased virus-cell attachment in a dose-dependent manner (**Figure 2F**). Notably, SLBR-N promoted higher levels of virus-cell attachment than SLBR-H or SLBR-B. This pattern closely mirrored the relative enhancement of JRFL infectivity observed in **Figure 2A**, supporting the conclusion that SLBR lectins augment virus infectivity by binding virions and promoting their attachment to target cells.

### Staphylococcal SSL lectins enhance HIV-1 infectivity in a strain- and lectin-dependent manner

We next evaluated the effect of SSL lectins on HIV infectivity. In the same DEAE dextran-free infectivity assay described above, treatment of JRFL and REJO.c with SSL3, SSL4, or SSL11 enhanced infectivity to varying levels in a lectin-and virus strain-dependent manner (**Figure 3A**). Mutations disrupting the sialic acid-binding motifs of these lectins (SSL3m, SSL4m, SSL11m) abolished infectivity enhancement, demonstrating the requirement for glycan recognition. The enhancing activity of SSL3 was reduced by sialylated O-glycan-rich mucin but not high mannose-N-glycan-bearing mannan (**Figure 3B**), confirming a sialoglycan-dependent mechanism. Similar to SLBRs, SSL lectins also interacted directly with HIV-1 virions and this binding was diminished by mutations at the sialic acid-binding site (**Figure 3C**). Among the three SSL lectins, SSL3 displayed the highest binding activity to both JRFL and REJO.c.

**Figure 3.**
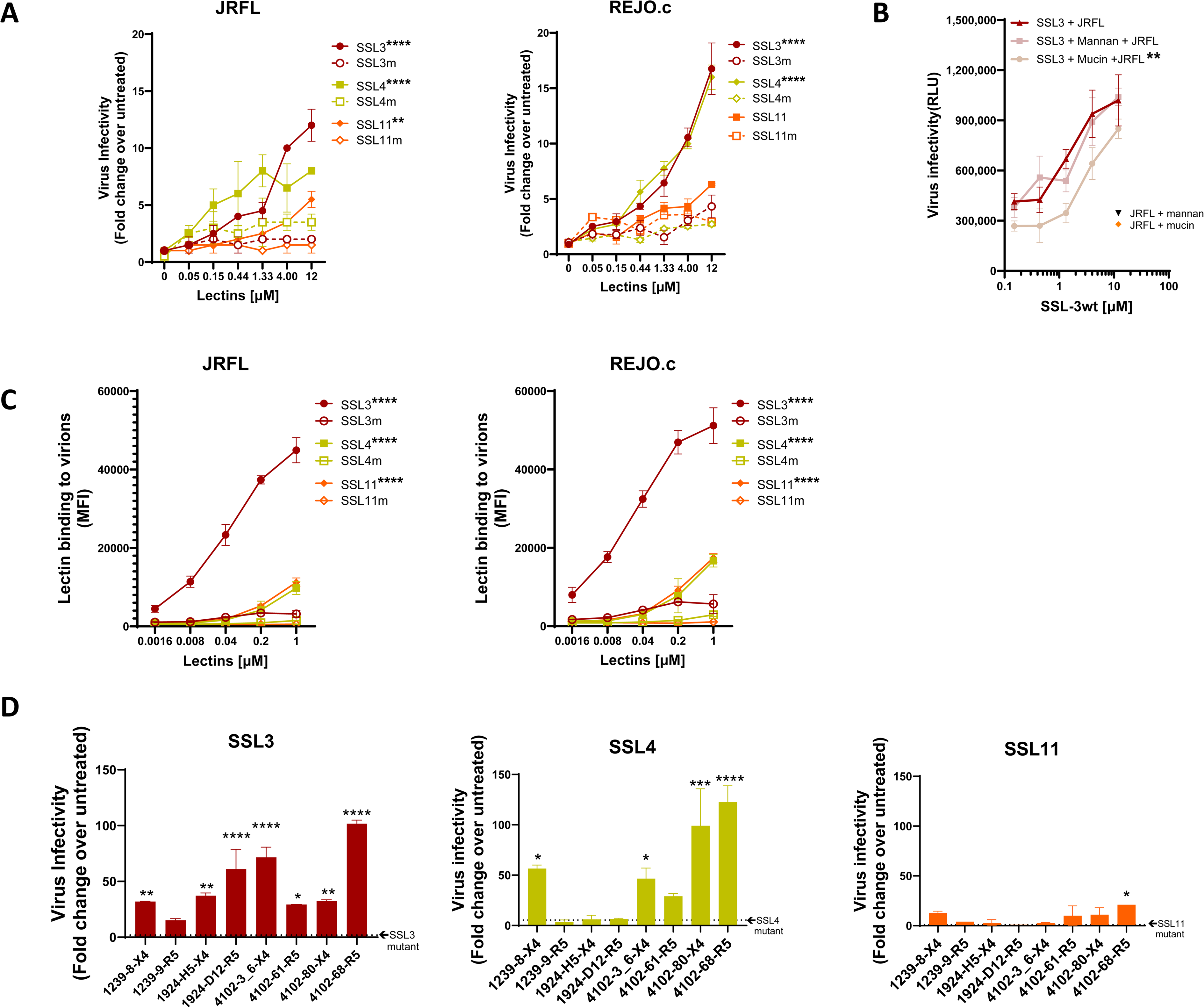
Staphylococcal superantigen-like (SSL) lectins SSL3, SSL4 and SSL11 enhance HIV-1 infectivity across different HIV-1 strains to varying degrees. **(A)** Enhanced infectivity of HIV-1 virions produced in 293T cells upon treatment with SSL lectins. Virions were pre-treated with titrated amounts of SSL lectins for 1 hour at 37°C and then incubated with TZM-bl reporter cells for 48 hours. Two clade B IMCs (JRFL and REJO.c) were tested with sLe^X^ -binding SSL lectins and their respective mutants. Untreated virions served as controls (set to 1). Infection with untreated JRFL and REJO.c yielded 10,855 RLU and 14,202 RLU, with a background (cell alone) level of 3,012 RLU and 2,895 RLU, respectively * p<0.05, *** p<0.001, **** p<0.0001 for comparison of each SSL wt vs. its respective mutant by two-way ANOVA with Sidak’s multiple comparison. **(B)** SSL3 enhancement activity sensitive to sialyated O-glycan-rich mucin and not high-mannose N-glycan-rich mannan. SSL3 was pre-incubated with 1 mg/mL mucin or mannan for 1 hour at 37°C and then incubated with JRFL for an additional 1 hour. Virus infectivity in TZM-bl reporter cells was measured 48 hours after infection. ** p<0.01 for comparison of JRFL + SSL3 treated with mucin vs. JRFL + untreated SSL3 by ANOVA with Kruskal-Wallis multiple comparison test. p >0.05 for JRFL + SLBR-N treated with mannan vs. JRFL + untreated SLBR-N. Orange diamond and black triangle: RLU for JRFL infection in the presence of only mucin or mannan, respectively. **(C)** SSL bound HIV-1 virions. Virus-SSL binding was measured using a Luminex bead assay, in which virions were coupled onto beads and treated with serial dilutions of His_6_-tagged SSL wt or mutant. SSL binding was then detected using biotinylated anti-His tag antibodies and PE-streptavidin. MFI: mean fluorescence intensity. **** p<0.0001,* p<0.05 for comparison of each SSL wt vs. its respective mutant by two-way ANOVA with Sidak’s multiple comparison. **(D)** Enhanced infectivity of X4-tropic and R5-tropic clade B IMCs produced in primary CD4 T cells upon treatment with SSL3, SSL4, and SSL11 lectins (each at 12 µM). ****p<0.0001 ***p<0.001 **p<0.01 *p<0.05 vs. no lectin by ANOVA. p≥0.05 is left unmarked. Data (mean ± SD) shown in each panel were from one experiment out of two or more repeat experiments.

The SSL lectin activity was also tested using eight X4 and R5-tropic clade B IMCs produced in activated PBMCs (59). SSL3, SSL4, and SSL11 heightened infectivity of seven, four, and one virus strains, respectively, again demonstrating substantial variability among virus strains (**Figure 3D**). Overall, SSL3 and SSL4 mediated greater enhancement than SSL11, although the magnitude of enhancement was lower than that observed with SLBR-N and SLBR-B (**Figure 2D**). For all SSL lectins tested, mutation of the sialic acid-binding site abrogated the enhancing activity, confirming the dependence on lectin-glycan interactions. Together with the SLBR data, these results show that distinct bacterial lectins that recognize sialylated glycans can enhance HIV infectivity in a glycan-dependent and virus-strain-specific manner. Moreover, the greater overall enhancement observed with SLBR vs. SSL lectins indicates that binding to O-linked sialoglycans can have a more profound effect than binding to N-linked sialoglycans.

### SLBR-N, but not SSL3, enhances HIV infection in primary CD4 T cells

Because prior studies (36) and experiments described above assessed the effects of bacterial lectins on HIV infection using TZM-bl reporter cells, we sought to examine whether similar effects were observed in primary target cells. HIV-1 reporter IMCs encoding heterologous *env* gene sequences from strains JRFL and C.1086 and expressing secreted Nano-luciferase (snNLuc) were pretreated with increasing concentrations of SLBR-N or SSL3 and then used to infect PHA-activated PBMCs or CD4 T cells obtained from two to three individual donors. Viral infection was quantified 48 h post-infection by nano-luciferase activity and expressed as fold change relative to untreated virus.

SLBR-N treatment significantly enhanced JRFL Env-mediated infection in PHA-activated PBMCs, with a consistent increase observed across donors (**Figure 4A top**). In contrast, SSL3 exhibited no detectable effect under the same conditions. A similar pattern was observed in activated CD4 T cells, where SLBR-N treatment induced a pronounced, dose-dependent enhancement of infection, while SSL3 again failed to increase virus replication across the concentrations tested (**Figure 4B top**). SLBR-N also increased C.1086 infection in PHA-activated PBMCs, although this activity was observed only with PBMC #6 and not with cells from the other donors (**Figure 4A-B, bottom**). SSL3 had no effect on C.1086 infection in all primary cells tested.

**Figure 4.**
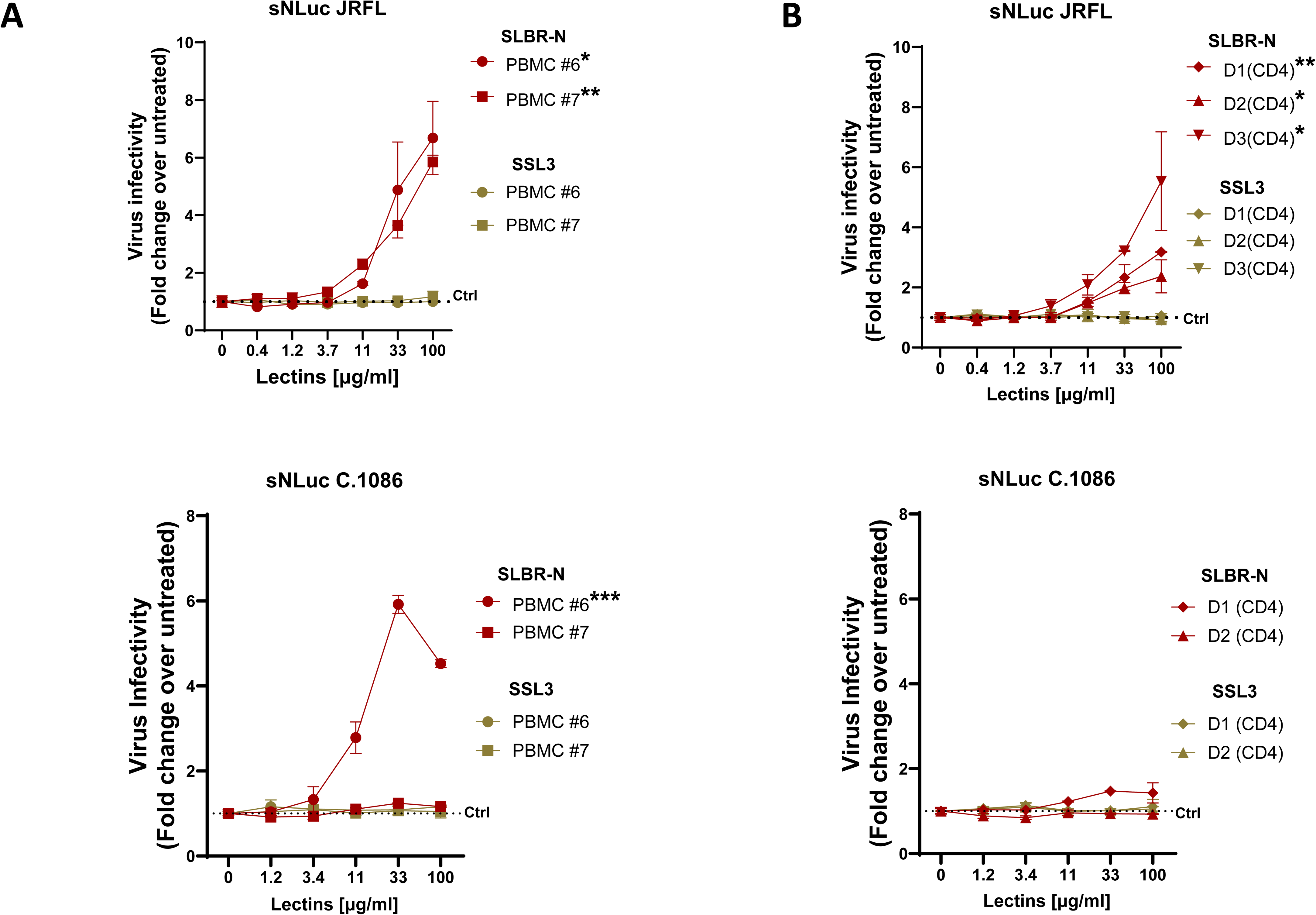
Effects of bacterial lectins SLBR-N and SSL3 on HIV-1 infection in primary CD4 T cells. JRFL and C.1086 IMC virions bearing a nanoluciferase (sNLuc) reporter gene, produced in 293T cells, were pre-incubated with titrated amounts of SLBR-N or SSL3 lectins for 1 hour at 37°C, and incubated with PHA-activated PBMCs (**A**) or enriched CD4 T cells (**B**) from different donors. Virus infection was measured after 48 hours using the Promega Nano-Glo Dual Luciferase Reporter Assay System. Experiments were performed in duplicate and repeated at least 3 times. Data from one of the experiments are shown as fold changes over untreated control (set to 1, dotted line). *p<0.05 **p<0.01 by two-way ANOVA comparing the effects of SLBR-N vs. SSL3 on JRFL or C.1086 infection in the same PBMC or CD4 T cell donors.

These results demonstrate that SLBR-N, but not SSL3, enhances HIV-1 infection in activated PBMCs and CD4 T cells, revealing lectin-specific differences in the ability of bacterial lectins to promote HIV infectivity in primary human T cell targets. The results also further support the importance of binding to O-linked rather than N-linked glycans for augmenting HIV infectivity.

### SLBR-N enhances HIV-1 trans-infection via DC-SIGN-independent and dependent pathways

We further evaluated whether SLBR-N modulates HIV trans-infection. We first examined virus trans-infection from bystander cells lacking the endogenous mannose-binding C-type lectin DC-SIGN. JRFL IMC-derived virions were pre-incubated with titrated concentrations of SLBR-N and then allowed to interact with either 293T cells or RAJI cells, both of which do not express DC-SIGN. After extensive washing to remove unbound virus, these cells were co-cultured with TZM-bl target cells, and trans-infection was quantified by beta-galactosidase activity 48 hours later. SLBR-N treatment increased HIV-1 trans-infection from both 293T and RAJI cells in a dose-dependent manner, with significant enhancement detected starting at 0.4 µM SLBR-N over untreated controls (**Figure 5A**).

**Figure 5:**
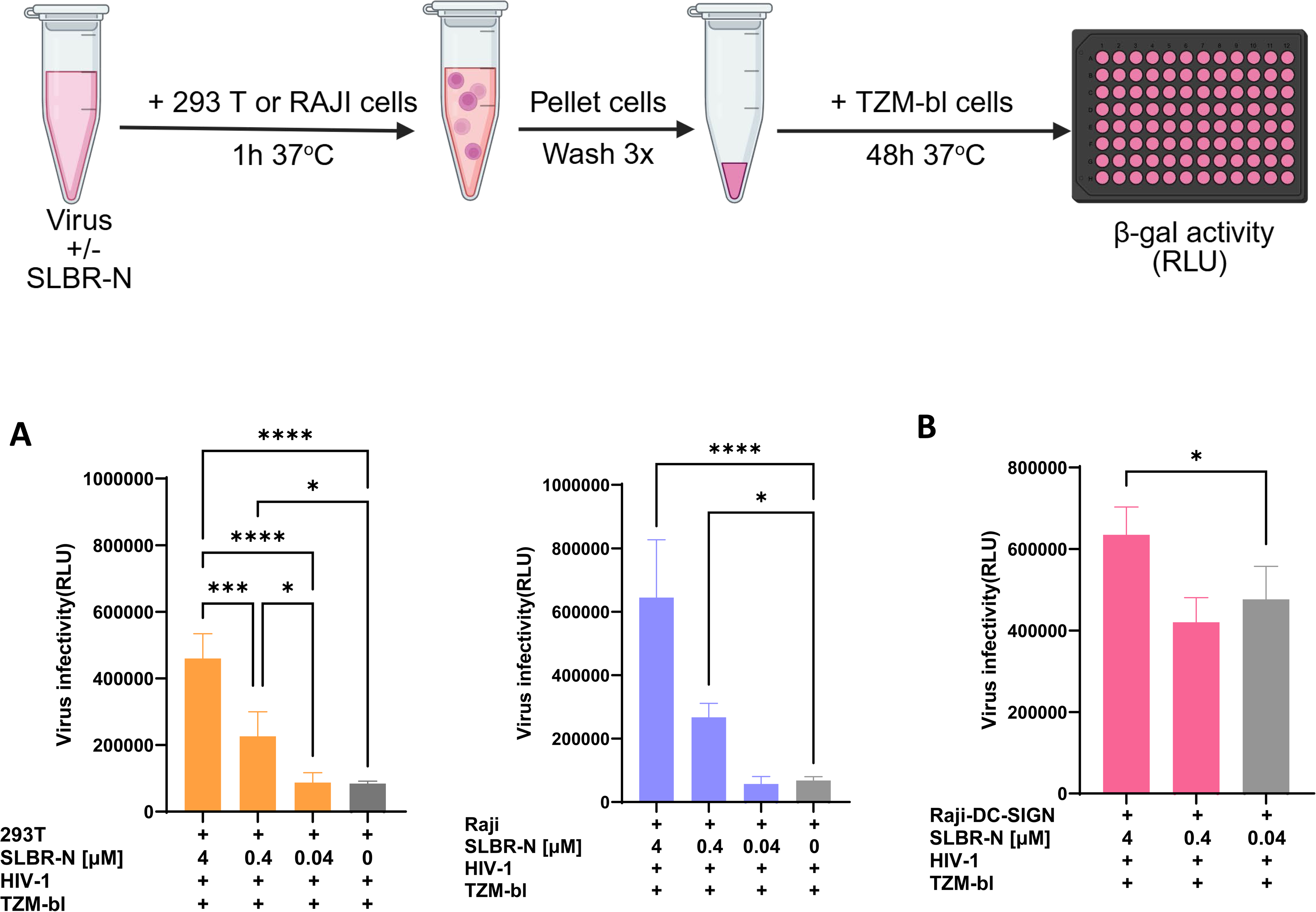
SLBR-N enhanced HIV-1 trans-infection via DC-SIGN-independent and dependent pathways. HIV-1 JRFL virions pre-treated with SLBR-N at varying concentrations were incubated with 293T and RAJI cells (A) or DC-SIGN^+^ RAJI cells (B) for 1 hour 37°C with periodic mixing. The cells were then extensively washed to remove unbound virions and co-cultured with TZM-bl cells for 48 hours 37°C. Luciferase activity (RLU) was detected as a measure of virus trans-infection to TZM-bl cells. * p<0.1, ***<0.001, **** p<0.0001 by one-way ANOVA with Tukey’s multiple comparisons. Experiments were performed in duplicate and repeated at least 3 times. Mean and standard error from the combined experiments are shown. p≥0.05, not marked.

We then assessed the effect of SLBR-N on trans-infection from DC-SIGN⁺ RAJI cells. As expected, we observed high baseline levels of HIV-1 trans-infection attributable to DC-SIGN alone. Addition of SLBR-N further elevated trans-infection, although the magnitude of enhancement was only ∼1.3-fold at the highest concentration tested (**Figure 5B**).

Collectively, these data indicate that SLBR-N can efficiently promote HIV-1 trans-infection in the absence of DC-SIGN, effectively substituting for DC-SIGN-mediated virus capture and transfer. The lack of inhibition of DC-SIGN-dependent trans-infection further suggests that SLBR-N does not interfere with DC-SIGN function, consistent with the distinct glycan specificities of these two lectins. These findings support a model in which SLBR-N enhances HIV-1 trans-infection through both DC-SIGN–independent mechanisms and auxiliary effects in DC-SIGN–expressing cells.

### SLBR-N enhances HIV-1 infection and virus burden in humanized mice

Finally, we determined whether SLBR-N influences HIV-1 infection in vivo using humanized NSG (NOD scid gamma) mice. Mice (n=9 per group) were derived from two independent cohorts engrafted with cells and tissues from different human donors and randomly assigned to the two groups. Mice were rectally inoculated with JRFL IMC that had been pretreated with SLBR-N or left untreated (**Figure 6A**). To model repeated mucosal exposure, mice received three sequential challenges with increasing virus doses (350, 700, and 1400 TCID₅₀). Plasma vRNA levels were monitored weekly by qRT-PCR.

**Figure 6:**
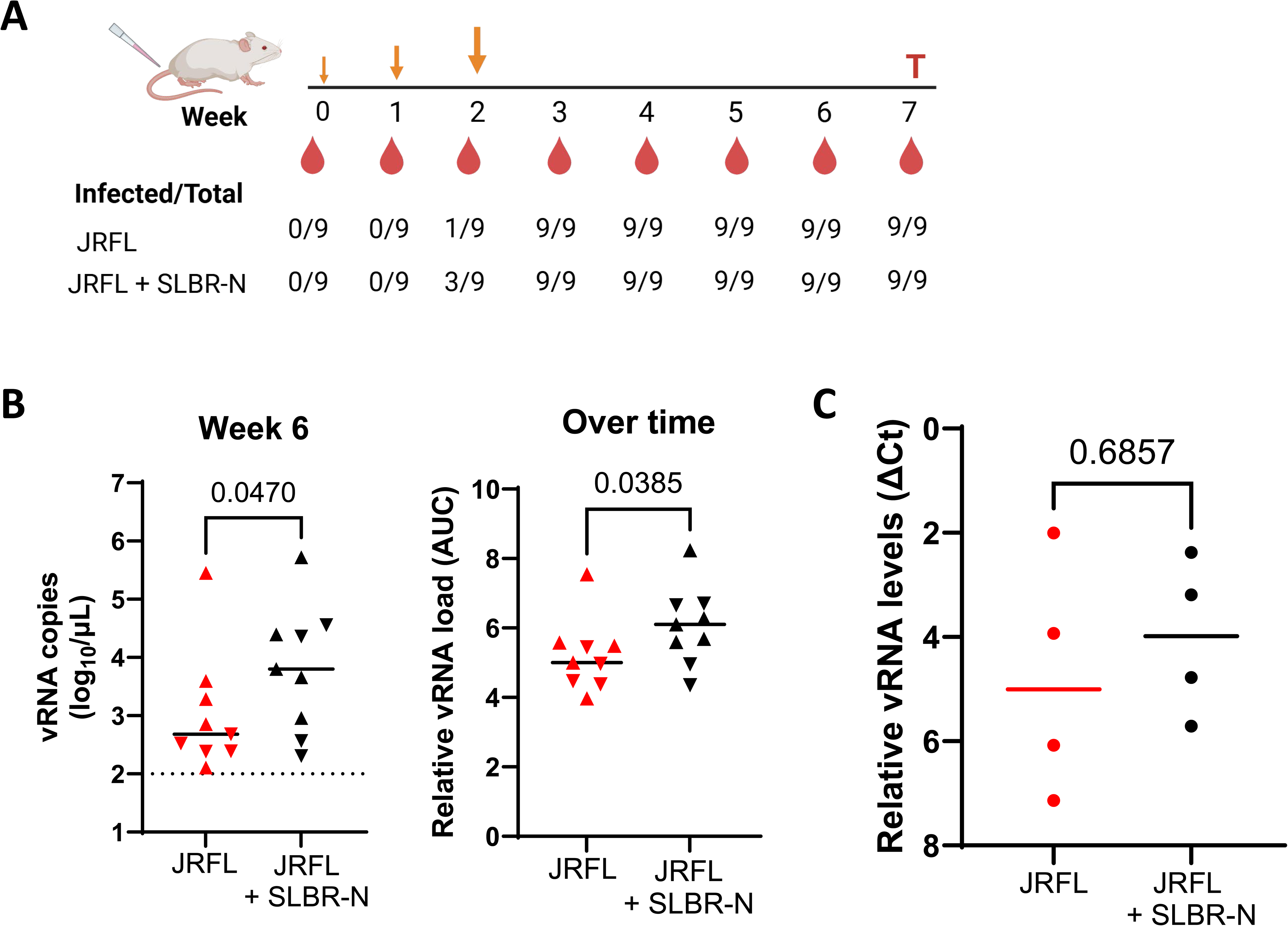
Virus burden in humanized mice after rectal inoculation of HIV-1 JRFL in the presence or absence of SLBR-N. **(A)** Experimental design. Virus was pretreated with or without SLBR-N for 1 hour 37°C and administered rectally to humanized mice (n=9) at three time points with increasing virus doses (350, 700, 1400 TCID50). Blood was collected weekly for vRNA quantification by RT-qPCR. The numbers of infected mice with vRNA copies above background are shown at the designated time points for the two groups. At the end of the study, spleen cells were collected to detect cell-associated vRNA. **(B)** vRNA load at week 6 and over time from week 2 to week 7 (as calculated by area under the curve, AUC). Two cohorts of mice generated from different human donors were assigned randomly to the two groups (up or down triangles, n=9/group). Dotted line: lower limit of detection. p values by Mann-Whitney test. **(C)** Relative vRNA levels in spleen cells from four mice from one of the cohorts at week 7 as measured by RT-qPCR. vRNA levels were calculated as ΔCt relative to GAPDH levels in the same samples. p values by Mann-Whitney test.

Two weeks after the first inoculation, plasma vRNA above background was detected in one mouse in the control group, compared with three mice in the SLBR-N-treated group (**Figure 6A**). Following subsequent virus challenges, from week 3 onward, all animals in both groups became viremic. Notably, mice exposed to SLBR-N-treated virus exhibited higher overall viral burdens than controls, as evidenced by both week-6 plasma viremia and longitudinal viral loads calculated as area under the curve (AUC) (**Figure 6B**). Analysis of cell-associated vRNA in splenocytes collected at the study termination (n=4 per group from the second cohort) further revealed a trend toward higher viral burden in mice receiving virus with SLBR-N compared with controls (**Figure 6C**). With the limited sample size, statistical significance was not achieved. Taken together, these results demonstrate that SLBR-N enhances mucosal HIV-1 infection and systemic viral burden in vivo in a humanized mouse model. These findings are in agreement with the in vitro data that SLBR-N mediates virion binding, promotes virus-cell attachment, and enhances virus infection and trans-infection across multiple cell types, supporting a role for bacterial lectins in modulating HIV-1 transmission and early infection dynamics.

## Discussion

In this study, we demonstrated for the first time using a humanized mouse model that the Streptococcal sialic-acid binding lectin SLBR-N facilitated mucosal HIV-1 infection in vivo, as indicated by increased initial transmission and higher virus burdens after infection was established. This activity is associated with the ability of SLBR-N to bind HIV-1 virions and promote virion attachment to target cells in a sialic acid-dependent manner, resulting in enhanced infectivity in both TZM-bl reporter cells and primary CD4 T cells. SLBR-N also promoted HIV-1 capture by bystander cells and increased virus trans-infection. Although SLBR-N further augmented DC-SIGN-mediated trans-infection, prominent effects were observed in the absence of DC-SIGN, suggesting that soluble SLBR-N may facilitate trans-infection by promoting virus interactions with bystander cells, independent of CD4, CCR5 coreceptor, and DC-SIGN, although the mechanisms have not been defined. Whether SLBR-N-bound virions are internalized into bystander cells and how they are transferred to target cells remains unknown. This observation is reminiscent of prior findings with soluble host lectins. For example, soluble sialic acid-binding Siglec-7 enhances HIV-1 entry into CD4⁺ T cells and macrophages (60). Galectin 8 also promotes HIV-1 and HIV-2 infection by acting as a molecular bridge between virions and target cells via its dual glycan-binding domains: an N-terminal domain recognizing sialyated glycans on virions and a C-terminal domain binding to terminal galactose on CD4 T cells (61). Together, findings presented here substantiate and extend our prior work (36), supporting a model in which HIV-1 virions can directly engage bacterial lectins at mucosal surfaces to enhance virus-cell attachment, direct and trans-infection of CD4 T cells, and ultimately transmission efficiency.

Similar to SLBR-N, other SLBR lectins (SLBR-H and SLBR-B) and staphylococcal SSL lectins from Staphylococcus aureus (SSL3, SSL4, and SSL11) also enhanced HIV-1 infectivity to varying extents, depending on the lectin and virus strain, although SLBR-N exhibited the greatest overall potency. The variability likely reflects both the fine specificity of each lectin and the sialoglycan heterogeneity across virus strains. Env (gp120/gp41) glycosylation is highly diverse across HIV-1 strains, with many unique N-and O-linked glycosites for each strain, each of which is decorated by an array of assorted glycan structures (56, 62, 63). Notably, changes in the Env signal sequence have been shown to alter Env glycan contents and thereby modulate Env interactions with host lectins, plant-derived antiviral lectins, and antibodies against glycan-bearing epitopes and glycan-dependent epitopes (56, 64). Consistent with this concept, we previously observed that SLBR-N reduced virus neutralization by broadly neutralizing antibodies targeting the V1V2 apex and the CD4-binding site (36), indicating that lectin binding exerts steric or allosteric changes in Env. In addition, lectin activity was dependent on target cells, as shown by disparate effects of SSL3 in TZM-bl cells versus primary CD4 T cells. The reasons are yet unknown and need further investigation.

It is important to note that enhancement of HIV-1 infectivity has thus far been restricted to sialic acid-binding lectins. In contrast, high mannose N-glycan-specific lectins such as E. coli FimH and lactobacillus Msl bind HIV-1 without increasing virus infectivity. SLBR and SSL lectins tested here are specific for sialoglycan structures, such as sLe^X^ and sLn, commonly found on N- and O-glycans. Although Env is particularly rich in N-glycans, with up to 30 N-glycans per Env protomer, O-glycans also have been detected by mass spectrometry and cryo-electron microscopy, particularly in the V1 region and at the conserved C5 site near the gp120/gp14 cleavage site (62, 63, 65). Computational prediction by NetOGlyc also identified potential *O*-glycan sites within the V1V2 and C5 regions (36). While we previously showed sialic acid-dependent Env reactivity with SLBR-N, the specific SLBR-binding sites remain undefined. Virus strain-specific lectin effects observed among X4 and R5-tropic viruses sharing identical backbones but distinct Env sequences implicate Env glycans as the primary lectin targets. However, HIV-1 virions also incorporate O-glycosylated host proteins, such as CD162 (P-selectin glycoprotein ligand-1), CD43 (sialophorin), and CD44 (E-selectin ligand) (66, 67), which may play a role in lectin binding, although their contributions remain unknown. The mechanisms by which sialic acid-binding lectins enhance HIV-1 infection are also undefined. The ability of these lectins to promote virus-cell attachment suggests that they may mask sialic acid residues on the virus surface and reduce electrostatic repulsion between the negatively charged virus and host cell membranes (68), similar to the effects of cationic polymers such as DEAE-dextran and polybrene (43, 45, 69), although it is yet unclear whether SLBR lectins can shield sufficient numbers of specific sialoglycans on the virus surface. In contrast, FimH and Msl lectins that bind to uncharged mannose residues do not exert similar effects (36).

Sialic acid-binding lectins are produced by multiple genera of mucosal bacteria. These lectins often function as adhesins or virulence factors that facilitate colonization and immune evasion. For example, the *S. gordonii* SLBR lectins are expressed on the tips of fimbriae to mediate attachment to receptors, such as salivary mucin MG2/MUC7 and platelet GP1ba that are implicated in oral colonization and endocarditis, respectively (38). The SabA adhesin of *Helicobacter pylori* with phylogenetic relationship with SLBR also binds to sialylated ligands (70). Similarly, *Mycoplasma genitalium* expresses a sialic acid-binding lectin called P110 or MgpC, which interacts with P140 and forms a transmembrane Nap complex on the outer bacterial surface to serve as adhesins to sialylated receptors on host cells (71). *M. genitalium*, which causes urethritis in men and cervicitis and pelvic inflammatory disease in women, has been associated with increased HIV-1 risk (72, 73). In comparison, Staphylococcal SSL lectins are secreted proteins that downmodulate immune responses by interacting with myeloid cells, inhibiting TLR activation, or preventing IgA binding to FcαRI on neutrophils (50–53). Despite these precedents, the prevalence and functional impact of sialoglycan-binding lectins within the mucosal microbiota remain largely unexplored.

These findings may have particular relevance in clinical scenarios in which medical interventions perturb the mucosal microbiota, potentially modulating HIV-1 risk. Genital secretions from Black and Hispanic women who received Depo Provera enhanced HIV-1 infectivity ex vivo (18, 74). In this population, Depo Provera administration was associated with disruption of the vaginal bacterial network, signified by loss of Lactobacillus connections and persistence of those with Streptococcus, Mycoplasma, and a limited number of other species (18). Given the documented association between Depo Provera use and increased risk of HIV-1 acquisition (13–17), these observations underscore the need to investigate further whether bacterial lectins contribute to the heightened virus susceptibility under such conditions.

## Conclusion

This study identifies sialic acid-binding lectins from common bacterial species of the human microbiota as potential modulators of HIV-1 transmission. These lectins interact directly with HIV-1 virions in a glycan-specific manner, promoting virus attachment to host cells and increasing virus infectivity through both direct infection and trans-infection pathways. Moreover, the presence of these lectins promotes HIV-1 infection in the context of mucosal transmission in a humanized mouse model. Collectively, these findings provide new insights into how the human microbiota may influence HIV-1 transmission across mucosal surfaces and underscore the importance of microbial-viral interactions as contributors to HIV-1 acquisition and pathogenesis.

## Limitations

This study primarily utilized recombinant bacterial lectins rather than native lectins expressed on streptococcal fimbriae and secreted by staphylococcal species. While this reductionist approach enabled attribution of virus infectivity-enhancing activity specifically to the lectins, further studies are needed to validate these effects in the context of intact bacteria within complex mucosal environments that include diverse host and microbial factors.

We also demonstrate, for the first time, that bacterial lectins can facilitate HIV-1 trans-infection independently of the host lectin DC-SIGN. Although the use of 293T and RAJI cell lines allowed clear discrimination of trans-infection mediated by bacterial lectins vs. DC-SIGN, these systems do not fully reflect physiologic conditions. Further studies should incorporate primary cell types, including vaginal, cervical, and foreskin epithelial cells, as well as dendritic cells expressing DC-SIGN, Siglecs, and other host lectins.

Finally, the in vivo experiments were conducted in a limited number of humanized mice. Although consistent results were observed with two different donor cohorts, this model has inherent limitations: it does not fully recapitulate the structural and functional characteristics of human mucosal tissues; its NSG background, which lacks B cell and antibody repertoires, precludes the establishment and maintenance of a representative human mucosal microbiota (75); and HIV-1 infection was achieved using high-dose rectal challenges that do not reflect the relatively low efficiency of sexual transmission in humans (76).

## Materials and methods

### Ethics statement

Animal work was reviewed and approved by the Icahn School of Medicine at Mount Sinai Institutional Animal Care and Use Committee.

### Cells, lectins, and viruses

HEK293T cells were purchased from the American Type Culture Collection (ATCC). The following reagent was obtained through BEI Resources, NIAID, NIH: TZM-bl Cells, HRP-8129, contributed by Dr. John C. Kappes, Dr. Xiaoyun Wu, and Tranzyme Inc (77). The following reagent was obtained through the NIH HIV Reagent Program, Division of AIDS, NIAID, NIH: parental Raji Cells (Raji-0), ARP-9944, and DC-SIGN-expressing Raji Cells (DC-SIGN+ Raji), ARP-9945, contributed by Dr. Li Wu and Dr. Vineet N. KewalRamani. These cell lines were maintained in Dulbecco’s modified Eagle medium (DMEM, Lonza) supplemented with 10% heat inactivated fetal bovine serum (FBS), penicillin/streptomycin (100U/mL), and L-glutamine.

PBMCs were isolated from Leukopaks purchased from the New York Blood Centre, and CD4^+^ T cells were purified from human peripheral blood using the MagCellect Human CD4^+^ T cell isolation kit (R&D Systems). Primary cells were maintained in complete RPMI 1640 medium supplemented with 10% FBS, 100U/mL penicillin, 100 μg/mL streptomycin, and 2mM L-glutamine.

SLBRs were expressed in *E. coli* transformed with the corresponding plasmids and purified as described (36, 48). SSLs were expressed in E. coli and purified as previously described (51).

Proviral IMC plasmids pNL-sNluc.6ATRi-B.JR-FL.ecto/K4760 and pNL-sNluc.6ATRi-C.1086.B2.ecto/K4830 encode *env* sequences of HIV-1 strains B.JRFL and C.1086.B2, respectively, and are engineered to express the secreted Nano-Luciferase (sNLuc) reporter (78, 79). The IMC plasmid of B.JRFL was obtained from Dr. Jerome A. Zack (UCLA) (80). The following reagents were obtained through BEI Resources, NIAID, NIH: Human Immunodeficiency Virus Type 1 (HIV-1) Infectious Molecular Clone, pREJO.c/2864, HRP-11746: pREJO.c/2864, contributed by Drs. John Kappes and Christina Ochsenbauer (55). These IMCs were generated by transfecting HEK293T cells using JetPRIME (Polyplus). Viral supernatants were filtered (0.45-micron) and pelleted through a 20% sucrose cushion by ultracentrifugation to enrich for virions while minimizing glycoconjugates from serum and cellular sources. Viral pellets were resuspended in PBS, titrated on TZM-bl cells, aliquoted, and stored at −80°C. Replication-competent HIV-1 clones 1239-8, 1239-9, 1924-H5, 1924-D12, 4102-3, 4102-61, 4102-80, and 4102-68 were produced in transfected 293T cells and then passaged on primary CD4 T cells (59).

### Virus infectivity assay

Virus infectivity in TZM-bl target cells was performed as described in (36), except that DEAE-Dextran was excluded and Beta-Glo® Assay System (Promega) reagents were used for detecting virus infection. Infection in primary peripheral mononuclear cells or CD4^+^ T cells was measured using the Nano-Glo® Luciferase Assay System (Promega). For these assays, virions were pre-incubated with titrated amounts of lectins for 1h at 37 °C prior to the addition of target cells (5000 cells/well) and incubated for 48h at 37°C. Each condition was tested in duplicates or triplicates. Assay controls included replicates of target cells alone (cell control), target cells with virus alone (virus control), target cells with lectin alone (lectin control). To test the effects of sialyated O-glycan-rich mucin (bovine submaxillary gland mucin, Thermo Scientific, Cat #: J63859MC) versus high-mannose-bearing mannan (Sigma-Aldrich, Cat#: M7504), lectin was pre-treated with each of these reagents (1mg/mL) and then incubated with virus.

### SLBR lectin binding to virions by glutathione beads

Virions (1 × 10⁸ vRNA copies/mL) was incubated with GST-tagged SLBR lectin (0.4 µM each) or left untreated, for 1 h at 37°C. Mucin-treated lectin was also tested to assess glycan-specific effect. Pierce^TM^ Glutathione Magnetic Agarose beads (100 µL; ThermoFisher) were then added and incubated for 1 h at room temperature. Beads were pelleted, washed three times with PBS to remove unbound virus, and quantified for bead-associated viral RNA by RT-qPCR using QIAamp Viral RNA Mini kit (Qiagen, Cat # 52904). cDNA was synthesized using the High-Capacity RNA to cDNA Kit (ThermoFisher, Cat # 4387406) and quantified by real-time PCR using the ABI PRISM 7900HT system. The assay was also performed after virus-lectin incubation for 24 h at 37°C and yielded comparable results (data not shown).

### Virus attachment to cells in the presence of lectin

Virions were pre-incubated with lectins at 37°C for 1h, then added to TZM-bl cells (1 × 10⁶ cells) on ice and maintained for another 1 h to allow binding, cells were then washed three times to remove unbound virions and lectins. vRNA on cells was quantified using RT-qPCR as described above.

### SSL lectin binding to virions on Luminex beads

SSL lectin binding to virions was quantified using a Luminex multiplex bead assay. Sucrose cushion-purified pre-titrated virions were covalently coupled to Luminex beads with the XMAP antibody coupling kit and incubated with titrated amounts of His-tagged SSL wt and mutant. Lectins bound to virions were detected using biotinylated anti-His antibodies (Abcam), followed by PE-labeled streptavidin (Abcam). Data were acquired with a Luminex Bio-Plex FlexMAP 3D system.

### Trans-infection

Virus trans-infection assay was performed as described in (81). JRFL IMC virions were pre-incubated with titrated amounts of SLBR-N for 1 hour at 37°C. Cells (HEK293T, Raji-0, RAJI-DC-SIGN) were added and incubated for an additional 1 hour at 37°C. After washing three times with culture medium to remove unbound virus, cells were co-cultured with TZM-bl reporter cells at a 1:1 ratio for 48 hours at 37°C. Infection of the TZM-bl cells was quantified using a luminescence-based β-galactosidase assay (Beta-Glo, Promega).

### Humanized mouse experiments

Humanized mice were purchased from the Columbia University Humanized Mouse Core (CUHMC, New York, NY). These mice were generated as previously described (82). Briefly, six- to eight-week-old female NSG mice received sublethal total body irradiation (1 Gy) using an X-ray irradiator (RS-2000, Rad Source Technologies, Suwanee, GA), followed by surgical implantation of a 1-mm³ human fetal thymic fragment under the kidney capsule. Mice also received intravenously human fetal liver hematopoietic stem cells (HSCs, 1 × 10⁵ cells) purified using a Human CD34 MicroBead Kit (Miltenyi Biotec, Cologne, Germany). Peripheral blood samples were collected by submandibular bleeding at 8 and 12 weeks post-transplantation, and human hematopoietic chimerism was analyzed by flow cytometry using fluorochrome-conjugated monoclonal antibodies (anti-mouse CD45-FITC and Ter119-PerCP-Cy5.5, as well as anti-human CD45-BV605, CD3-AF700, CD4-BV786, CD8-PE/Dazzle594, CD19-PE, and CD14-APC/Cy7 (all from BioLegend, San Diego, CA). Flow cytometric analysis was performed using an LSRFortessa (BD Biosciences, San Jose, CA), and data were analyzed with FlowJo software (Tree Star, Ashland, OR).

Mice with >20% human CD45+ leucocytes reconstitution were selected for experiments and randomly assigned into two groups (n=5/group). Mice were rectally inoculated with increasing doses of JRFL IMC (350, 700, or 1400 TCID50) administered either with or without SLBR-N (0.06 μM in 10 μL/animal). Blood was collected periodically for measurement of plasma viral load.

### Measurement of plasma vRNA and cell-associated vRNA

vRNA was extracted from plasma using the QIAamp viral RNA Mini kit, while total RNA from blood and tissue was extracted using the Qiagen RNeasy Mini kit. cDNA was prepared using a High-Capacity cDNA Reverse Transcription kit with RNase Inhibitor (ThermoFisher, Cat # 4374966) on a C1000 Touch Thermal Cycler. HIV cDNA was then quantified by real-time PCR using a Taqman-based assay on an ABI 7500 Fast Real Time PCR System (Applied Biosystems) as previously described (83). The HIV *gag* sequences of the forward and reverse primers and the Taqman probe were 5’-CAG CCA AAA CTC TTG CTT TAT GG-3’, 5’-GGG ACC AGC AGC TAC ACT AGAA-3’, and NDP:5’-6FAM TGA TGA CAG CAT GCC AGG GAG TGG MGBNFQ-3’, respectively. Cell-associated vRNA was measured simultaneously with cell-derived GAPDH RNA, which was quantified using a Pre-developed Taqman assay reagents kit (Applied Biosystems, Cat # 4333746F).

### Statistical analyses

Data analyses were performed using GraphPad Prism software using statistical tests specified in figure legends.

## Acknowledgments

This study was supported in part by the National Institutes of Health (grants AI150909, AI139290, and AI148327 to C.E.H., NIDCR R03DE029516 to B.A.B., AI042853, and AI172843 to M.S), the Department of Veterans Affairs (Merit review grant I01BX005616 to C.E.H. and Research Career Scientist Award IK6BX004607 to C.E.H.), the Gates Foundation (Grant ID: OPP1032144 to C.O.), and the Health Research Council of New Zealand (HRC 22/322 to R.J.L.)

We thank the staff members at the Icahn School of Medicine at Mount Sinai Center for Comparative Medicine and Surgery for animal husbandry and veterinary care, Dr. Hui Wang (Humanized Mouse Core at Columbia University Center for Translational Immunology) for generating humanized mice, Dr. Jeromine Klingler (Icahn School of Medicine at Mount Sinai) for preparation of PBMCs and CD4 T cells, Drs. Wei Chao and Nada Marjanovic (Icahn School of Medicine at Mount Sinai) for assistance with RT-qPCR, and the late Dr. Jie Zheng (University of Alabama at Birmingham) who cloned proviral IMC plasmids pNL-sNluc.6ATRi-B.JR-FL.ecto/K4760 and pNL-sNluc.6ATRi-C.1086.B2.ecto/K4830.

## Author Contributions

CKY: Investigation, Methodology, Formal analysis, Visualization, Writing – original draft, writing review & editing

XL: Investigation, Methodology, Data curation, Visualization, Writing, - review & editing

RJL: Resources, Writing – review & editing

CO: Resources, Writing – review & editing

BAB: Resources, Writing – review & editing

FA: Resources, writing, review & editing

MS: Resources, Writing – review & editing

CEH: Conceptualization, Funding Acquisition, Project administration, Resources, Supervision, Writing – original draft, writing – review & editing

---

**Supplemental Figure S1.**
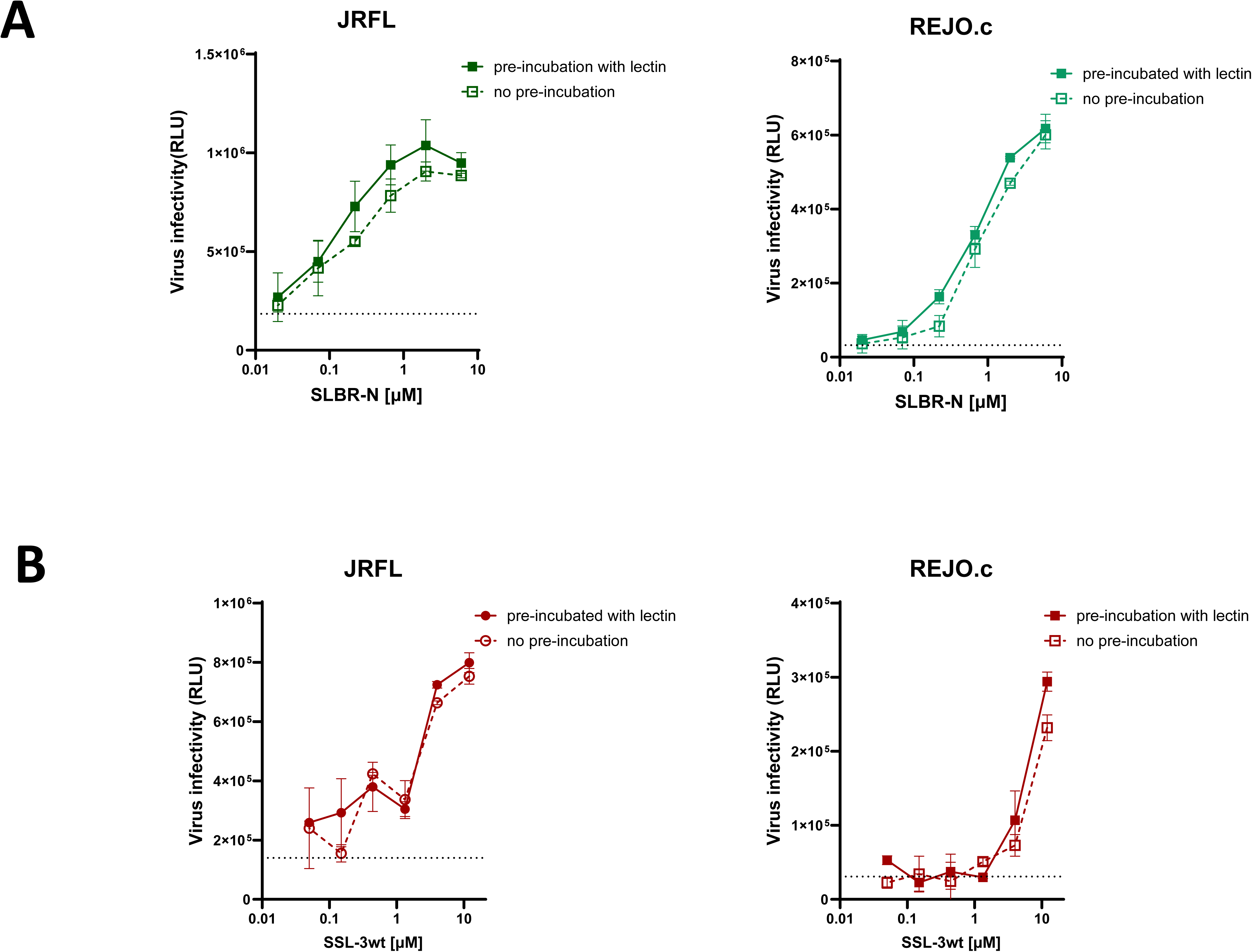
SLBR-N and SSL3 lectins enhanced HIV-1 infectivity with or without lectin-virus preincubation. JRFL and REJO.c IMC virions were either pre-incubated with SLBR-N (**A**) or SSL3 (**B**) for 1 hour at 37°C or added simultaneously to lectin and TZM-bl target cells without pre-incubation. Virus infectivity in TZM-bl reporter cells was quantified 48 hours post-infection. Mean ± SD are shown. RLU: relative luminescence unit. Dotted lines: virus infection without lectin.

**Supplemental Figure S2.**
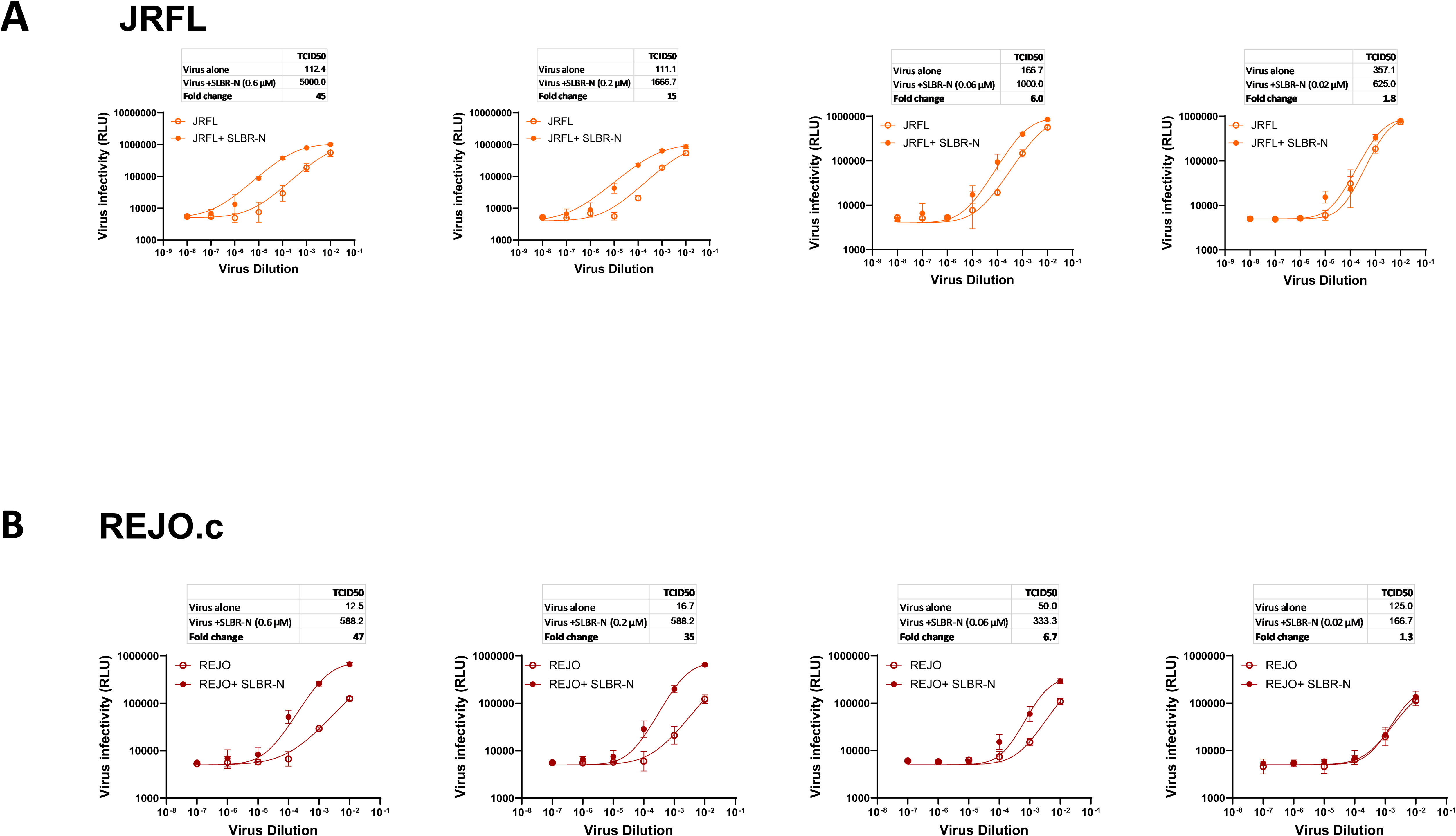
SLBR-N increases virus infectivity as measured by TCID50. Virions of JRFL (**A**) and REJO.c (**B**) IMC were serially diluted (10^−2^ to 10^−7^ or 10^−8^), treated with SLBR-N (0.6, 0.2, 0.06, and 0.02 µM) or left untreated at 37°C for 1 hour and then incubated with TZM-bl reporter cells for 48 hours. Virus infection was measured by beta-galactosidase activity. Reciprocal virus dilutions to attain 50% infection (TCID50) were calculated in GraphPad Prism using a non-linear fit model. RLU: relative luminescence unit.

